## Supplementary figures and images for "Dual STDP processes at Purkinje cells contribute to distinct improvements in accuracy and vigor of saccadic eye movements"

### Supplementary Fig 1

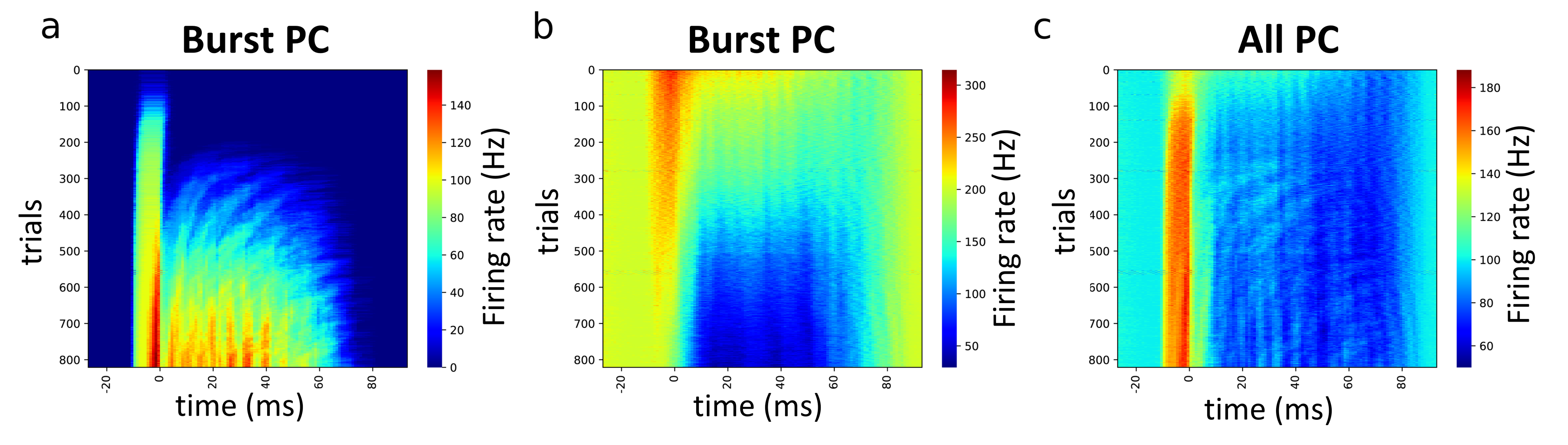

### Supplementary Fig 2

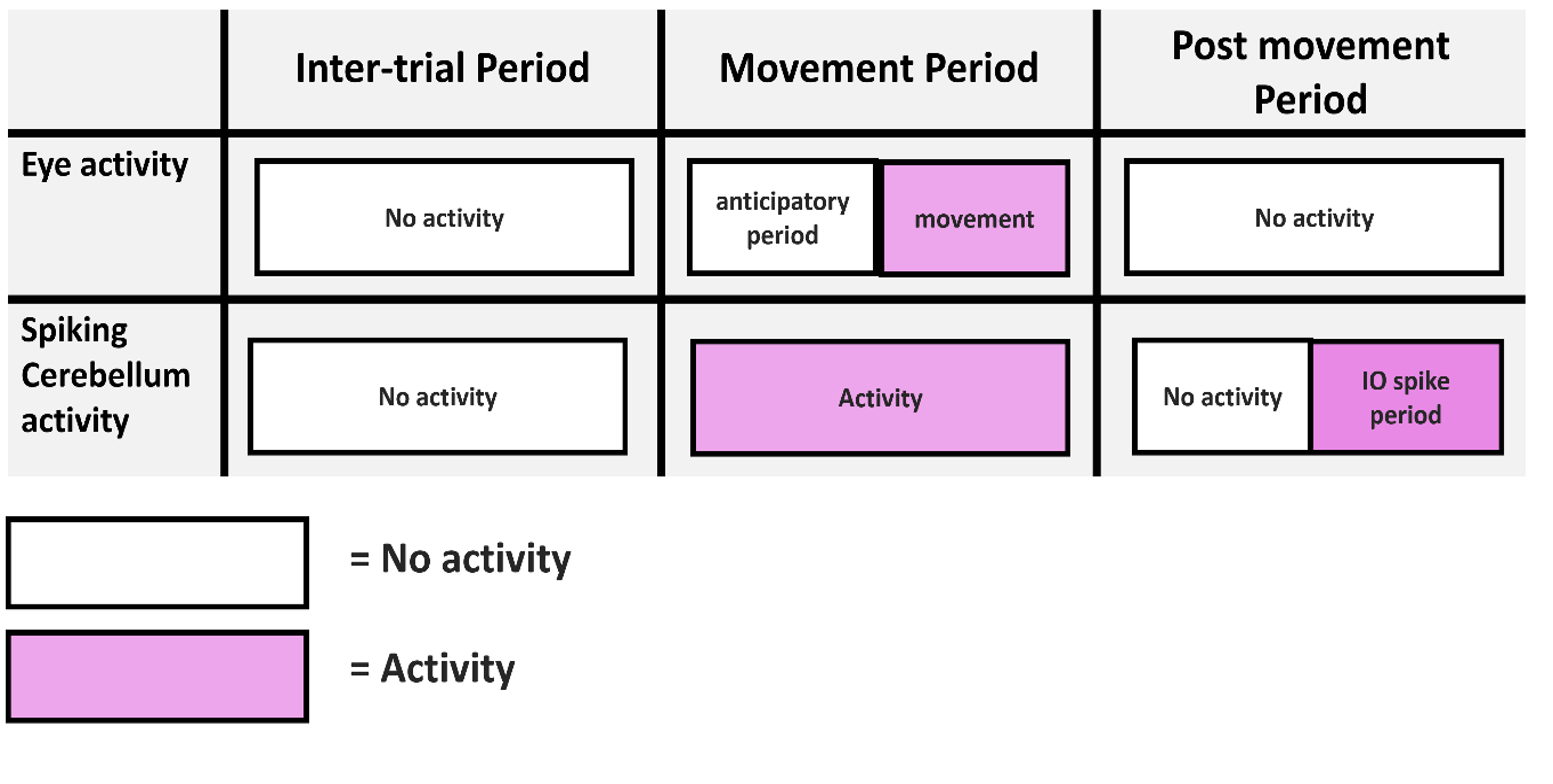
